# The α-synuclein hereditary mutation E46K unlocks a more stable, pathogenic fibril structure

**DOI:** 10.1101/868869

**Authors:** David R. Boyer, Binsen Li, Chuanqi Sun, Weijia Fan, Kang Zhou, Michael P. Hughes, Michael R. Sawaya, Lin Jiang, David S. Eisenberg

**Affiliations:** Departments of Chemistry and Biochemistry and Biological Chemistry, UCLA-DOE Institute, Molecular Biology Institute, and Howard Hughes Medical Institute, UCLA, Los Angeles, CA 90095; Department of Neurology, David Geffen School of Medicine, Molecular Biology Institute, UCLA, Los Angeles, CA 90095 USA; California Nano Systems Institute, UCLA, Los Angeles, CA 90095 USA; Department of Cell & Molecular Biology, St. Jude Children’s Research Hospital, Memphis, TN 38105 USA

**Author notes:** Correspondence to: Lin Jiang,; David S. Eisenberg. These authors contributed equally to this work (David R. Boyer, Binsen Li).

## Abstract

Aggregation of α-synuclein is a defining molecular feature of Parkinson’s disease, Lewy Body Dementia, and Multiple Systems Atrophy. Hereditary mutations in α-synuclein are linked to both Parkinson’s disease and Lewy Body Dementia; in particular, patients bearing the E46K disease mutation manifest a clinical picture of parkinsonism and Lewy Body Dementia, and E46K creates more pathogenic fibrils *in vitro.* Understanding the effect of these hereditary mutations on α-synuclein fibril structure is fundamental to α-synuclein biology. We therefore determined the cryoEM structure of α-synuclein fibrils containing the hereditary E46K mutation. The 2.5 Å structure reveals a symmetric double protofilament in which the molecules adopt a vastly re-arranged, lower energy fold compared to wild-type fibrils. We propose that the E46K misfolding pathway avoids electrostatic repulsion between K46 and K80, a residue pair which forms the E46-K80 salt-bridge in the wild-type fibril structure. We hypothesize that under our conditions the wild-type fold does not reach this deeper energy well of the E46K fold because the E46-K80 salt bridge diverts α-synuclein into a kinetic trap – a shallower, more accessible energy minimum. The E46K mutation apparently unlocks a more stable and pathogenic fibril structure.

**Significance Statement:** Parkinson’s is the second most prevalent neurodegenerative condition, leading to movement disorders, and dementia in some cases. Because of the strong association of this condition with amyloid aggregates of the protein α-synuclein, structural understanding of these amyloid aggregates may be the path to eventual therapies. Our study of the structure of a variant α-synuclein inherited in families afflicted with a clinical picture of parkinsonism and Lewy Body Dementia supplements recent structures of the wild type structure, and shows how a single residue change can result in a greatly changed structure that may underlie the inherited form of the disease.

## Introduction

The group of diseases termed the synucleinopathies - Parkinson’s Disease (PD), Lewy Body Dementia (LBD), and Multiple Systems Atrophy (MSA) – is thought to be caused by the aggregation of α-synuclein (α-syn) into amyloid fibrils. The causal relationship between the formation of amyloid fibrils of α-syn and the synucleinopathies is supported by several observations. Aggregated α-syn is a major component of Lewy Bodies, the hallmark lesion in PD and LBD, and the hallmark lesions of MSA^1, 2^. Hereditary mutations in α-syn are linked to familial forms of PD and LBD^3^. Overexpression of wild-type α-syn via dominantly inherited duplications and triplications of the gene that encodes α-syn, SNCA, are sufficient to cause PD^4–6^. Further, the injection of fibrils of α-syn into the brains of mice induced PD-like pathology including Lewy body and Lewy neurite formation, cell-to-cell spreading of Lewy body pathology, and motor deficits similar to PD^8^. Although it is never fully possible to establish causation, these combined observations suggest the case is strong for the linkage of aggregated α-syn to the synucleinopathies.

To gain a molecular level understanding of amyloid fibrils of α-syn, we previously applied cryo-EM to determine the near-atomic structures of fibrils of recombinantly assembled α-syn^7^. We observed two distinct structures – termed the rod and the twister – that share a similar structural kernel formed by residues 50-77 but differ in their protofilament interfaces and flanking regions of the fibril core (Figure 1 b, Figure 4 b-c, Supplementary Figure 1 a-b). The rod and twister structures display amyloid polymorphism – the formation of distinct fibril conformations by the same protein sequence.

**Figure 1.**
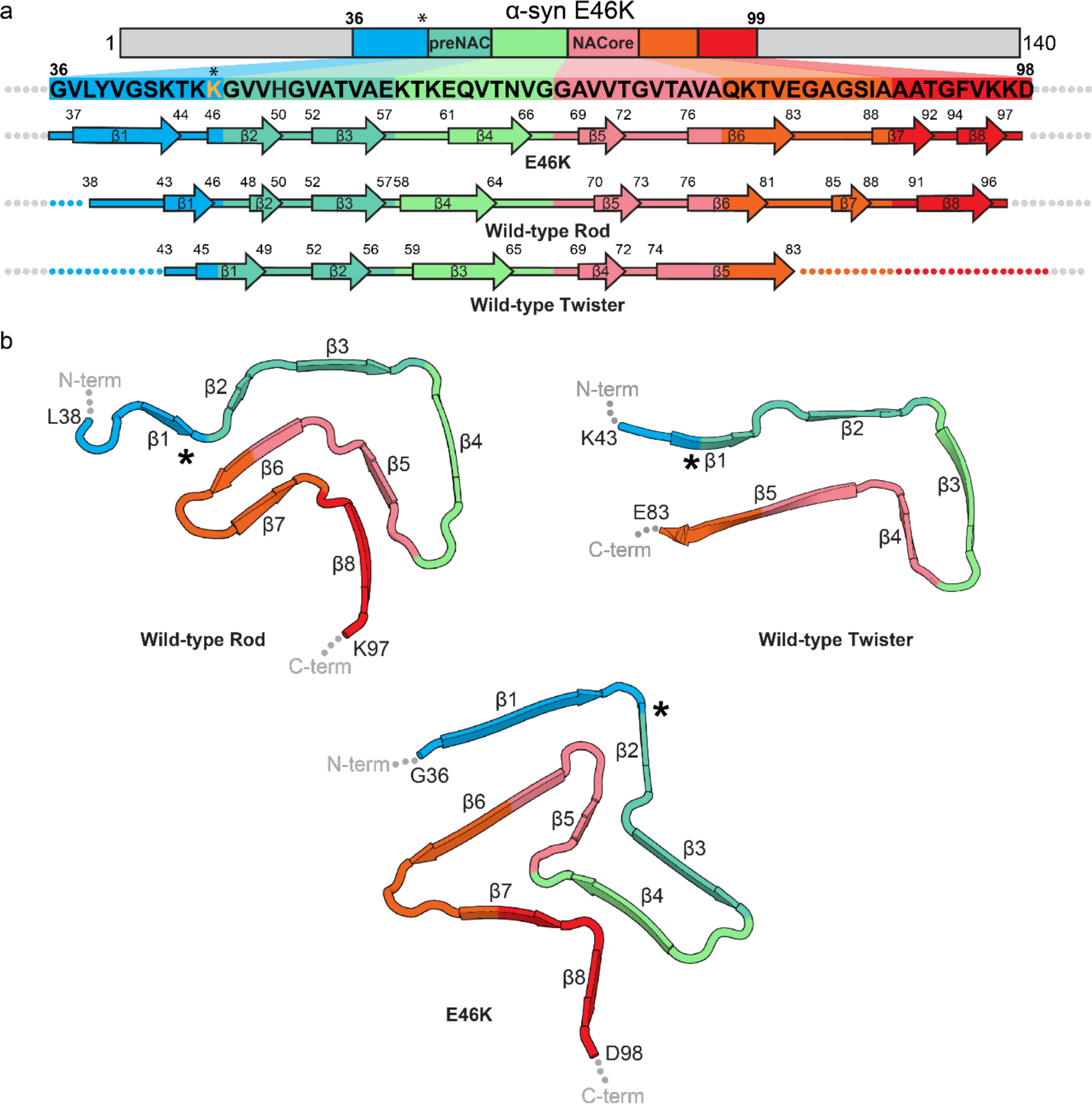
The E46K mutation leads to a re-packed protofilament fold a) Primary and secondary structure of α-syn wild-type and E46K fibrils. Arrows indicate regions adopting β-strand conformation while other residues form loops and turns. The E46K mutation lies N-terminal to the preNAC region (residues 47-56). b) Protofilament folds of the previously determined wild-type folds (rod and twister) and E46K protofilament fold determined here^7^. Asterisks indicate the location of residue 46.

Amyloid polymorphism has been observed in numerous other amyloid-forming proteins including the tau protein and amyloid-β^8–11^. For tau protein, the different polymorphs observed correspond to distinct diseases, namely Alzheimer’s, Pick’s disease, and chronic traumatic encephalopathy^8–10^. Although the structures of α-syn fibrils derived from human disease brain tissue have not yet been determined, previous research has shown that fibrils derived from PD have different biochemical properties than fibrils from MSA, including differential seeding activity and cell-type origin and infectivity, that may result from the formation of distinct polymorphs in the two diseases^12^. Therefore, atomic structures of fibril polymorphs are key to both basic understanding of amyloid protein structure and the development of disease-specific therapeutics.

Although hereditary mutations offer a crucial link between α-syn and disease, it is unknown how they might exert their effects. One hypothesis is that hereditary mutations may encode new structures of α-syn with enhanced pathogenicity. We note that new structures formed as the result of hereditary mutations are not true polymorphs – different structures adopted by the same protein sequence. Instead, we label them as “quasi-polymorphs” – different structures adopted by a protein pair with almost identical sequences.

Several hereditary mutations in α-syn may result in the formation of quasi-polymorphs. Mutations A30P, E46K, H50Q, G51D, A53E, and A53T have all been discovered to result in autosomal dominant synucleinopathies^3^. Of these, E46K seems to be the only hereditary mutation that manifests in a clinical picture closer to LDB whereas other mutations are found in the context of PD, suggesting that E46K may have a unique effect on the structure of amyloid fibrils of α-syn^13^. Indeed, solid state NMR studies of α-syn E46K show large chemical shift differences relative to wild-type fibrils, suggesting large scale rearrangements in the fibril structure as a result of the E46K mutation^14^.

Consistent with the evidence that E46K alters α-syn fibril structure and disease manifestation, E46K has been shown to increase the pathogenicity of α-syn fibrils compared to wild-type. *In vitro* studies have shown that E46K results in an increase in α-syn’s phospholipid binding ability and an enhancement of fibril formation^15^. In addition, E46K promotes higher levels of aggregation in cultured cells relative to wild-type, A53T, and A30P α-syn^16^. Further, others have found that α-syn bearing E46K is more toxic to rat primary neurons compared to wild-type, A30P, and A53T^17^.

Three previous structures of full-length wild-type non-acetylated as well as three structures of acetylated full-length and C-terminally truncated α-syn reveal that E46 participates in a conserved salt-bridge with K80 (Supplementary Figure 1 a)^7, 18–21^. The E46K mutation must eliminate this salt bridge due to electrostatic repulsion that disfavors the proximity of K46 to K80, potentially leading to a different fibril structure (quasi-polymorph). A structural difference such as this may help explain the altered biochemical properties and ssNMR chemical shifts of E46K α-syn fibrils. Therefore, we sought to determine the structure of α-syn fibrils containing the E46K hereditary mutation. We find that the E46K mutation produces a homogenous sample composed of a single species whose structure differs radically from structures determined thus far. Consistent with prior studies demonstrating increased pathogenicity of E46K α-syn, we also find that E46K fibrils are more powerful seeds in α-syn HEK293T biosensor cells and more strongly impair mitochondrial activity in PC12 cells. Combining structural, energetic, and biochemical analysis, we attempt to understand the structure-function relationship of α-syn fibrils.

## Results

### Cryo-EM structure determination and architecture of E46K α-syn fibrils

We purified α-syn bearing the E46K hereditary mutation and generated amyloid fibrils using the same growth conditions as in our previous studies^7^. We then optimized cryo-EM grids of E46K fibrils and imaged them at 165,000x magnification on an energy-filtered Titan Krios equipped with a K2 Summit direct electron detector operating in super-resolution mode. We also took advantage of image shift induced-beam tilt correction in SerialEM during data collection to maintain coma-free alignment during data collection^22^. Using helical refinement procedures in Relion 3.0, we obtained a 2.5 Å resolution reconstruction of E46K α-syn fibrils (Figure 2 a-e, Supplementary Figure 2 a-c, Supplementary Figure 3, Table 1).

**Figure 2.**
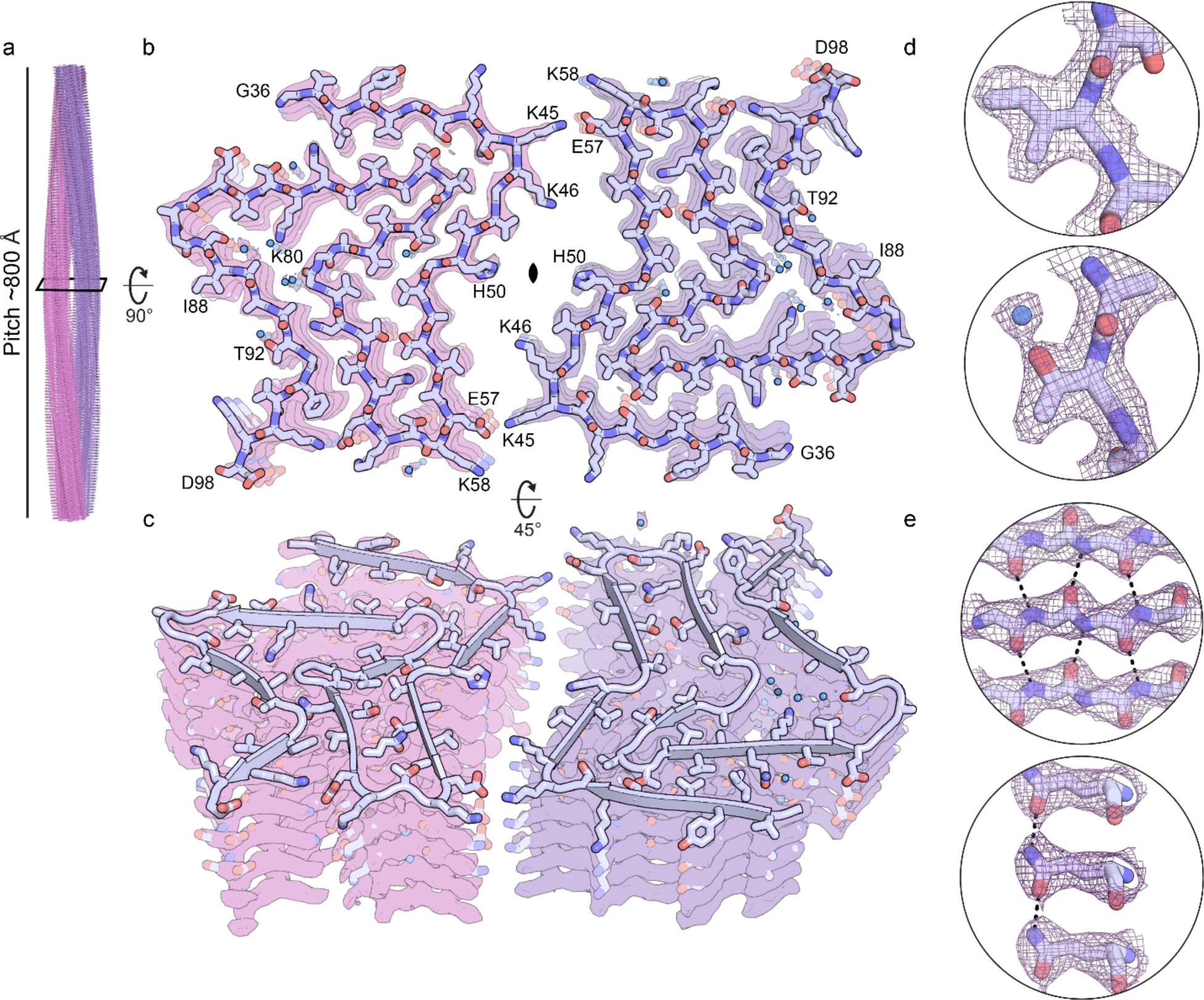
Overview of E46K cryo-EM structure a) E46K fibril side-view demonstrating a pitch of ∼800 Å. b) Cross-sectional view of one layer of the E46K fibril. The fibril is wound from two identical protofilaments related by a two-fold rotational symmetry axis. The E46K protofilament contains ordered residues 36-98. c) Tilted view of E46K fibril cross-section demonstrating the stacking of identical layers. d-e) Representative side chain and backbone densities highlight resolution of reconstruction and good map to model agreement. d) Coulombic potential map of Ile88 and Thr92. An ordered water molecule is bound to the oxygen of the threonine side chain. e) Coulombic potential map of representative backbone (Gln79-Val82) and side chain amide hydrogen bonding (Gln62).

**Table 1.**
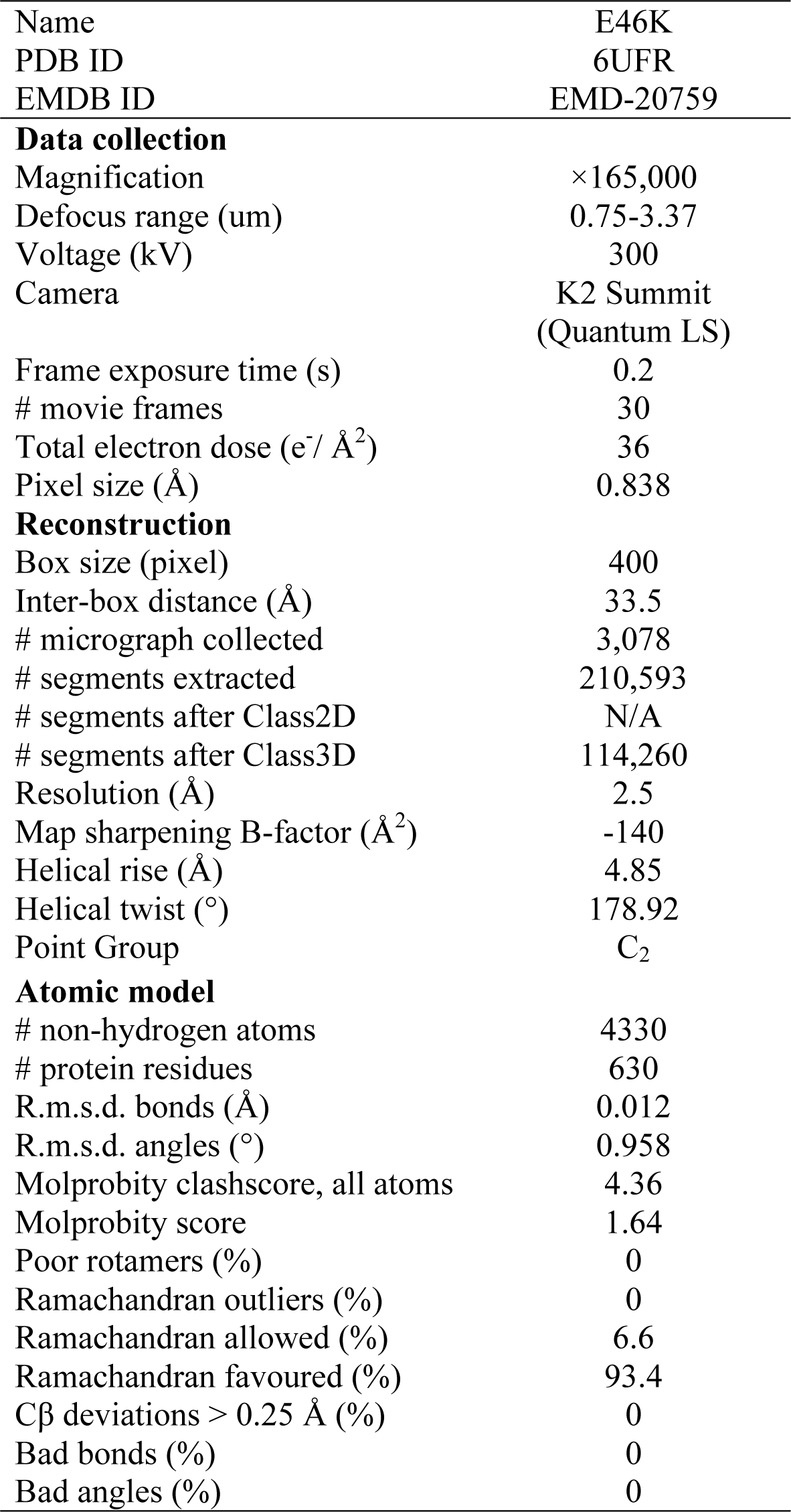
Cryo-EM data collection, refinement, and validation statistics. Information for number of segments after Class2D is not available as only Class3D was performed for 400 pixel box segments (see *Methods*).

The high resolution allowed us to confidently assign side chain rotamer and carbonyl positions, confirming the left handedness and the internal C_2_ symmetry of the helical reconstruction, and to build an unambiguous atomic model *de novo* (Figure 2 b-e, Supplementary Figure 3, see *Methods*). We were also able to visualize water molecules hydrogen bonded to polar and charged side chains such as threonine, serine, and lysine, as well as to backbone carbonyl oxygen atoms (Figure 2 d-e, Supplementary Figure 3). The atomic model reveals that the E46K fibrils are wound from two protofilaments formed by ordered residues Gly36 to Asp98 that come together at a “wet” two-fold symmetric interface formed by residues K45-E57 (Figure 2 b-c, Figure 3 a).

**Figure 3.**
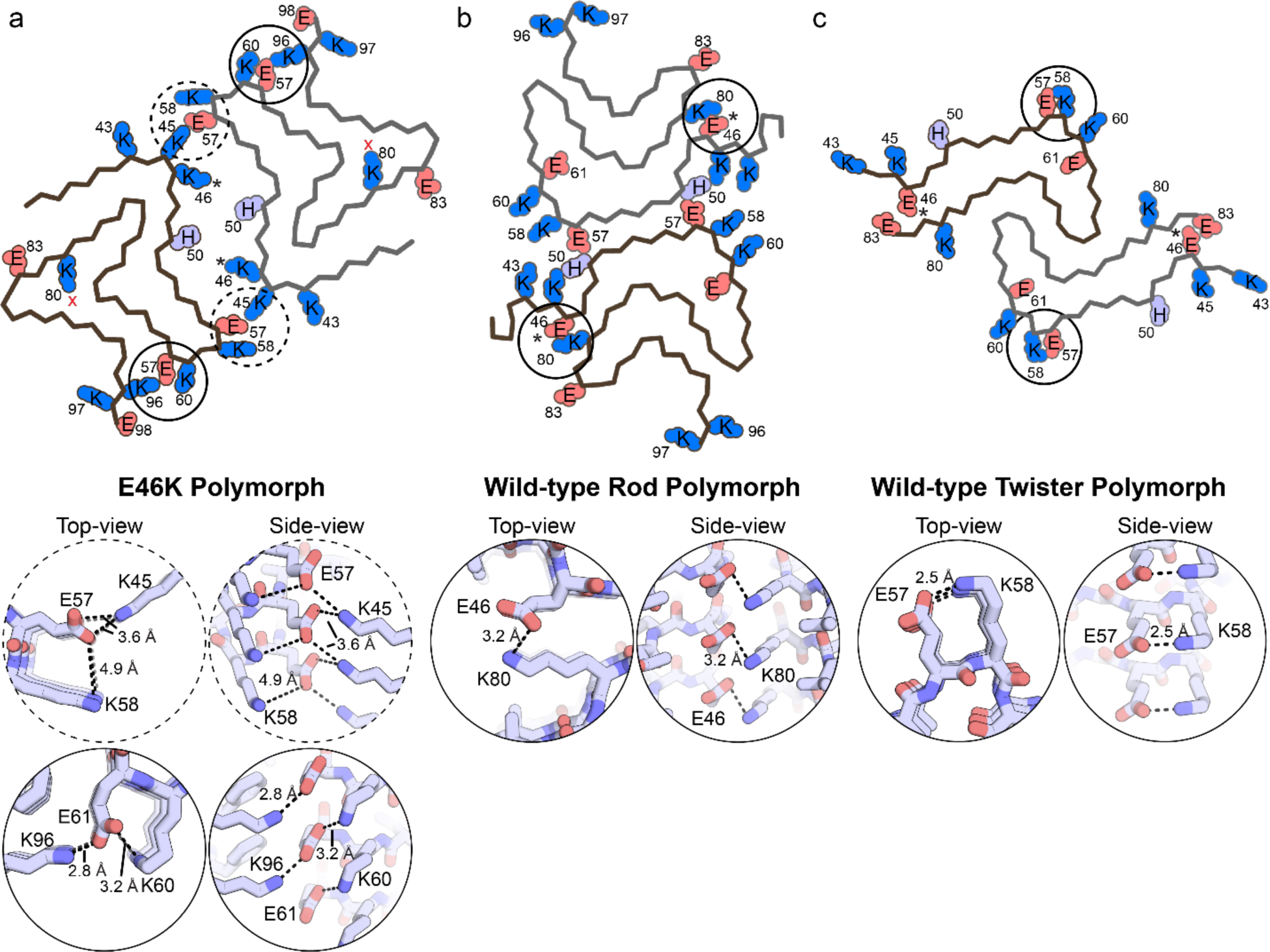
Electrostatic residues and interactions of wild-type and E46K fibrils a) Overview of E46K quasi-polymorph fold with charged and ionizable residues shown (top). Top- and side-view of the K45-E57-K58 and K60-E61-K96 electrostatic triads (bottom). The E46K fibril core has a net charge of +11 per layer (2*8 lysine/per chain + 2*1/2 histidine per chain – 2*3 glutamate per chain) and 8 charged pairs per layer (2*K45-E57 + 2*E57-E58 + 2*K60-E61 + 2*E61-K96). b) Overview of the wild-type rod polymorph fold with charged and ionizable residues shown (top). Top- and side-view of E46-K80 salt bridge (bottom). The wild-type rod fibril core has a net charge of +7 per layer (2*7 lysine/per chain + 2*1/2 histidine per chain – 2*4 glutamate per chain) and two charge pairs per layer (2*E46-K80 per chain). c) Overview of wild-type twister polymorph fold with charged and ionizable residues shown (top). Top- and side-view of E57-K80 salt bridge (bottom). The wild-type twister fibril core has a net charge of +3 per layer (2*5 lysine/per chain + 2*1/2 histidine per chain – 2*4 glutamate per chain) and two charge pairs per layer (2*E57-K58 per chain). Asterisks denote the location of residue 46. Red x’s denote location of the ordered water molecules hydrogen bonding to K80 in the E46K structure.

### Comparison of E46K and wild-type α-syn protofilament folds

Overall, the E46K mutation leads to a different protofilament fold than the previously observed wild-type rod and twister structures, resulting in a new quasi-polymorph of α-syn. Figure 1 a-b compares the secondary and tertiary structure of the E46K fibril with the wild-type rod and twister. Both the rod and twister form a similar structural kernel comprised of residues 50-77, in which β-strands β3, β4, and β5 in the rod and β2, β3, and β4 in the twister form a bent β-arch (Figure 2). The twister has fewer ordered residues at the N- and C-termini of its fibril core compared to the rod (Figure 1 a-b, Supplementary Figure 1). The C-terminal residues in the rod structure form a Greek-key like fold comprising β5, β6, β7, and β8. However, in the E46K structure, the kernel and Greek-key are not maintained and a new packing arrangement is formed. This is likely due to the E46K mutation disrupting the wild-type E46-K80 salt bridge, allowing a rearrangement of the backbone (Figure 2 b). Interestingly, K80 is now buried in the fibril core, although we visualize an ordered molecule binding to the primary amine of the K80 side chain, suggesting that an extensive hydrogen bond network may help to counteract the apparent unfavorable placement of this polar, charged residue within the fibril core. Without the constraint of the E46-K80 salt bridge, a new set of residues form sheet-sheet interfaces within the protofilament (Figure 2). Now tightly mated heterozippers are formed by β-strands β1 and β6, β3 and β4, and a roughly triangular-shaped bent β-arch fold is formed by β4, β5, β6, and β7 (Figure 1).

Supplementary Figure 4 a-c helps to visualize the changes in interacting residues among the structures by displaying all pairwise interactions found within the protofilaments of wild-type and E46K quasi-polymorphs. This analysis of pairwise interactions not only codifies the differences in sets of interacting residues between the wild-type and E46K structures but also clearly reveals that the E46K structure has more interacting residues than its wild-type counterparts. Correspondingly, each chain within the E46K protofilament has a greater buried surface area than wild-type structures (7,944 Å^2^ for E46K; 7,605 Å^2^ for wild-type rod; 5,082 Å^2^ for wild-type twister). In line with this observation, energetic analysis indicates that the E46K structure has a lower standard free energy (greater stabilization) than wild-type structures (Supplementary Figure 1 a-c, Table 2). To verify the energy estimate, we performed an SDS denaturation assay where both wild-type and E46K fibrils were incubated with various concentrations of SDS at 37 °C followed by ThT fluorescence measurements. Our results demonstrate that E46K fibrils are more resistant to chemical denaturation than wild-type fibrils, consistent with our energetic calculation suggesting that E46K fibrils are more stable than wild-type fibrils (Supplementary Figure 5 b).

**Table 2.** Comparative solvation energies

| Fibril Structure | Atomic solvation standard free energy of stabilization (kcal/mol) |  |  |
| --- | --- | --- | --- |
|  | Per Layer | Per Chain | Per Residue |
| E46K | -73.8 | -36.9 | -0.59 |
| WT CryoEM Rod (6CU7) | -59.6 | -29.3 | -0.49 |
| WT CryoEM Twister (6CU8) | -25.0 | -12.5 | -0.30 |
| Tau PHF (530L) | -62.8 | -31.4 | -0.43 |
| Tau Pick's (6GX5) | -44.8 | -44.8 | -0.48 |
| Tau CTE Type I (6NWP) | -67.4 | -33.7 | -0.48 |
| TDP-43 SegA-sym (6N37) | -34.2 | -17.1 | -0.47 |
| Serum Amyloid A (6MST) | -68.8 | -34.4 | -0.65 |
| FUS (5W3N) | -12.2 | -12.2 | -0.20 |

A key difference between E46K and wild-type structures is their pattern of electrostatic interactions. The E46K fibril core has a greater number of charged pairs (interacting pairs of glutamate and lysine) creating a more balanced set of electrostatic interactions despite having a higher net charge than the wild-type fibrils (Figure 3 a-c). This results from the E46K structure containing four electrostatic triads per layer featuring residues K45-E57-K58 and K60-E61-K96 while the wild-type rod and twister structures both contain only two electrostatic zippers featuring residues E46-K80 and E57-K58, respectively (Figure 3 a-c). Therefore, in line with the observation above that the E46K structure has overall more inter-residue contacts, a higher buried surface area, and a lower free energy than wild-type, it also has a richer set of electrostatic interactions.

Interestingly, although the wild-type and E46K structures differ, many residues adopt similar secondary structures. For instance, β-strand interrupting loops and turns are formed by KTKE pseudo-repeats (residues 43-46, 58-61, 80-83) and glycines (residues 51, 67-68, 73, 84, and 93) while other residues form β-strands (Figure 1 a-b). Indeed, if one compares the structures in Figure 1 b, especially the wild-type rod and E46K, loops and turns between β-strands often lead to a similar change in chain direction. For example, turns between β3 and β4 (right-turn), β4 and β5 (right-turn), β5 and β6 (left-turn), β6 and β7 (left-turn), β7 and β8 (right-turn) change the chain in generally similar manners. However, there are differences in the extent and radii of the turns that generate the structural diversity seen in the structures. For instance, the turn between β3 and β4 leads to a ∼90° turn in the rod and a ∼180° turn in the E46K structure. Also, the turn between β4 and β5 leads to a ∼180° turn in the rod and a ∼100° turn in the E46K structure. These observations highlight the critical importance of turn and loop regions in generating amyloid polymorphism.

### Electrostatic zippers constitute the E46K protofilament interface

Despite having a tighter protofilament fold that buries more surface area, the interface between E46K protofilaments contains fewer contacts than those in the wild-type rod and twister structures (Figure 4 a-c). Instead of the two protofilaments meeting at a classical steric zipper interface in which beta-sheets from each protofilament tightly mate with interdigitating side chains excluding water, the E46K structure forms a largely solvent-filled interface spanning residues 45-57 (buried surface area = 47.3 Å^2^). This is in contrast to the dry, steric zipper-like interfaces formed by the preNAC (buried surface area = 91.7 Å^2^) and NAC (buried surface area = 65.3 Å^2^) residues in the wild-type rod and twister, respectively. Although most residues are too far apart to interact in the E46K interface, two electrostatic zippers form on either side of the interface (Figure 4 a). Electrostatic zippers have previously been observed to enable counterion-induced DNA condensation whereby anionic phosphates and cations such as Mn^2+^ and spermidines alternate along the length of the DNA-DNA interface forming a “zipper” that “fastens” the molecules together^23^. In the E46K protofilament interface, electrostatic zippers consist of carboxylate anions of E57 interleaving with the K45 side chain aminium cations, fastening the two protofilaments together (Figure 4 a). This interaction is not only repeated twice per protofilament interface due to the two-fold helical symmetry, but extends for thousands of layers along the fibril axis. We note that the staggered nature of the electrostatic zipper leaves unpaired charges on both the top and bottom of the fibril, potentially attracting additional monomers to add to the fibril through long-range electrostatic interactions. We also observe that no single protofilaments were detected during class averaging indicating that, despite its relative lack of interactions across the protofilament, the E46K interface is apparently strong enough to consistently bind together the two protofilaments of the fibril.

**Figure 4.**
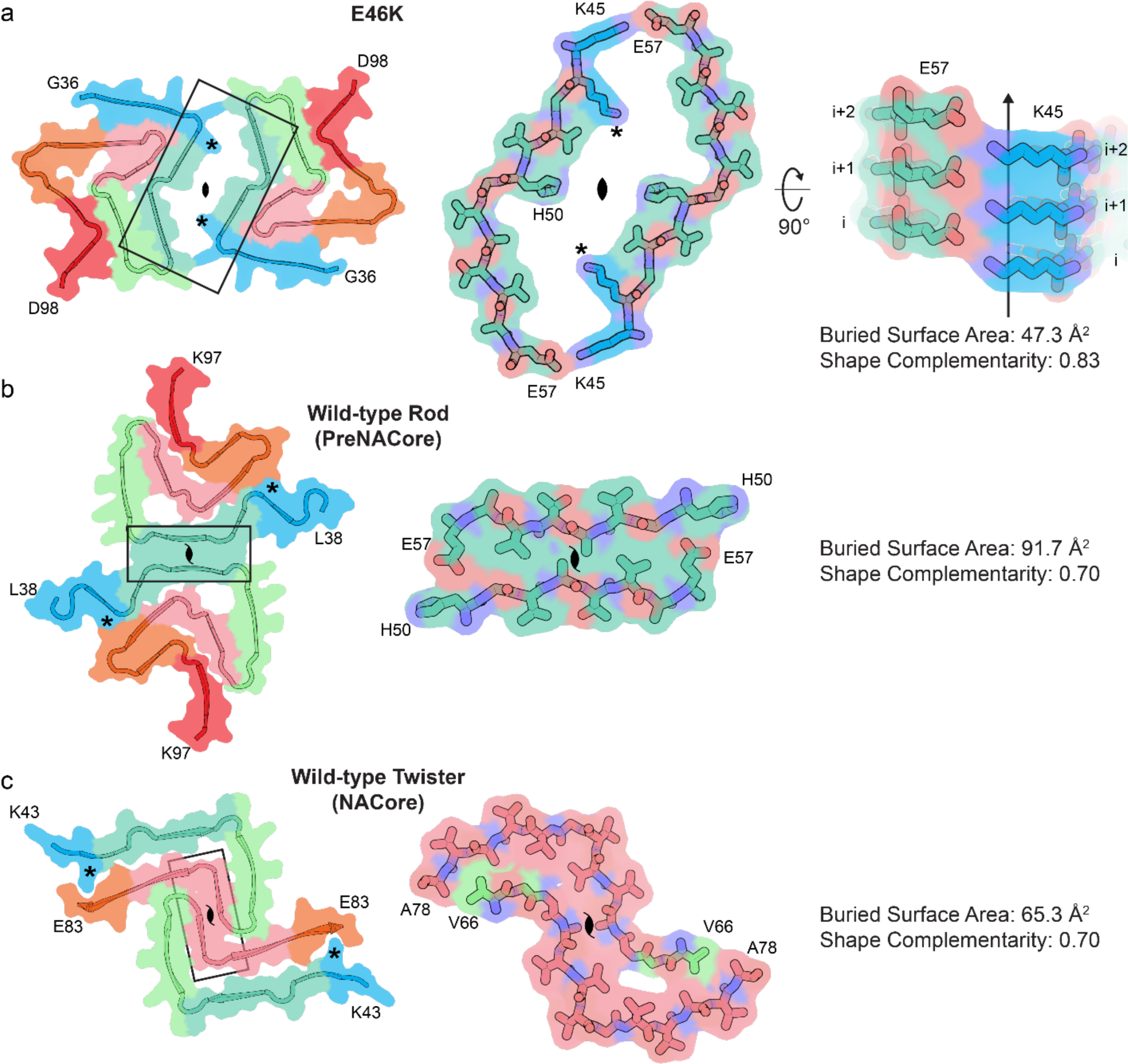
Comparison of protofilament interfaces of E46K and wild-type fibrils a) The E46K quasi-polymorph forms a two-fold symmetric protofilament interface comprised of residues Lys45-Glu57. The interface is largely solvent-filled and is held together by electrostatic interaction of Lys45 and Glu57 at both ends of the interface. The Lys45-Glu57 salt bridge is repeated along the length of the fibril creating an electrostatic zipper. b) The wild-type rod polymorph forms a pseudo-2_1_ symmetric dry steric zipper interface comprised of residues His50-E57 from the preNAC region. c) The wild-type twister polymorph forms a pseudo-2_1_ symmetric dry steric zipper interface comprised of residues Val66-Ala78 from the NACore.

### E46K fibrils are more pathogenic than wild-type

We wondered if the differences in structure between the E46K and wild-type fibrils resulted in differences in biochemical properties. Therefore, we first examined the ability of sonicated E46K and wild-type fibrils to seed endogenously expressed α-syn-A53T-YFP in HEK293T biosensor cells^24^. If seeding occurs, normally diffuse α-syn-A53T-YFP will aggregate into discrete puncta, which can be counted and used as a robust measure of seeding. Our results indicate that at most concentrations tested, E46K fibrils are significantly more powerful seeds than wild-type fibrils (Figure 5 a). Whereas the biosensor cell assay supports that E46K fibrils are better seeds than wild-type fibrils, the biosensor cells express α-syn-A53T-YFP, and not wild-type α-syn. In addition, differences in lipofectamine uptake may influence which fibril species is able to induce more seeding in the biosensor cell assay. Therefore, in order to directly examine whether E46K fibrils can seed wild-type protein more strongly than wild-type fibrils, we next tested the seeding ability of E46K fibrils in an *in vitro* assay (Supplementary Figure 5 a). Similar to the biosensor cell assay, E46K fibrils were more efficient than wild-type fibrils in seeding growth of wild-type α-syn (Supplementary Figure 5 a). Further, the different ThT binding ability of wild-type fibrils either grown without seeds or with wild-type seeds compared to wild-type fibrils seeded by E46K fibrils suggests that seeding by E46K fibrils can induce a different wild-type fibril structure, possibly one similar to the E46K fibril structure (Supplementary Figure 5 a). This is similar to what is seen in Watson *et al.* where both non-acetylated and acetylated α-syn form fibrils to the same extent, yet acetylated α-syn has a substantially lower ThT signal^25^. We also observe that unseeded E46K has a longer lag phase than unseeded wild-type α-syn; we speculate why E46K may form fibrils more slowly than wild-type in the Discussion.

**Figure 5.**
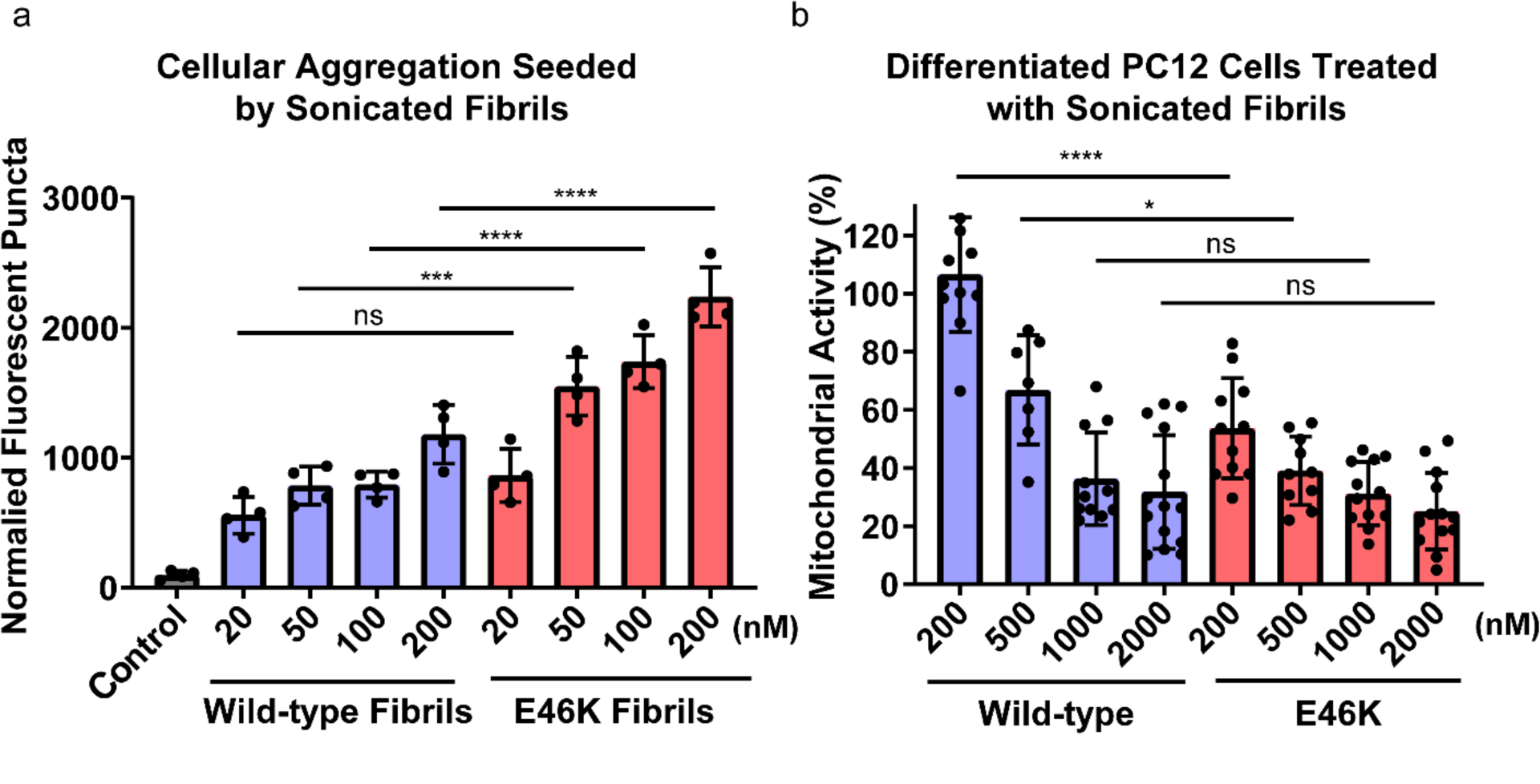
Biochemical analysis of E46K vs. wild-type fibrils a) Fibrils of E46K and wild-type α-syn were sonicated and transfected into HEK293T α-syn-A53T-YFP biosensor cells, and aggregation is measured by the normally soluble and diffuse intracellular α-syn-A53T-YFP forming discrete puncta, which are then quantified through fluorescent image analysis^24^. At all concentrations but 20 nM, E46K fibrils are significantly more powerful seeds than wild-type fibrils. Error bars represent the standard deviation of four technical replicates. b) In order to assay the toxicity of E46K and wild-type fibrils, we treated PC12 cells with sonicated fibrils and measured mitochondrial activity with an MTT assay^26, 27^. E46K fibrils require a lower concentration to significantly impair mitochondrial activity compared to wild-type fibrils. Error bars represent the standard deviation of 7-14 technical replicates. **** = p-value ≤ 0.0001. ** = p-value ≤ 0.01. ns = p-value > 0.05. P-values were calculated using an unpaired, two-tailed t-test with a 95% CI.

We next compared the ability of E46K and wild-type fibrils to impair mitochondrial activity of differentiated neuron-like rat pheochromocytoma (PC12) cells as a proxy for comparing cytotoxicities^26, 27^. We observe significantly impaired mitochondrial activity by lower concentrations of E46K fibrils than of wild-type fibrils (Figure 5 b). Consistent with previous results showing increased aggregation in SH-SY5Y cultured cells, and higher toxicity to rat primary neurons, our results indicate that the structure formed by E46K fibrils is more pathogenic than wild-type^15–17^.

## Discussion

### Structural differences help explain enhanced pathogenicity of E46K α-syn fibrils

The structural differences between E46K and wild-type fibrils help rationalize the differences in biochemical properties we observe. First, we have shown an increased seeding efficiency of E46K fibrils compared to wild-type in HEK293T biosensor cells and *in vitro* (Figure 5 a, Supplementary Figure 5 a). This difference may arise from the stronger electrostatic templating mechanism of E46K fibrils than wild-type fibrils: there are four electrostatic triads per layer in the E46K fibril structure compared to two E46-K80 salt bridges in the wild-type rod structure and two E57-K80 salt bridges in the wild-type twister structure, and each one of them forms a staggered electrostatic zipper, with overhanging, unsatisfied charges (Figure 3 a, c). Since all of these electrostatic residues are present in the α-syn-A53T-YFP construct in the biosensor cells and in wild-type α-syn used in the *in vitro* assay, it is plausible that the electrostatic triads in the E46K fibril guide α-syn monomers to the ends of E46K fibrils through long-range electrostatic attraction. Second, the E46K fibril has a different set of ordered surfaces than wild-type fibrils, which may lead to a different set of interacting partners in the cellular milieu. It has previously been shown that aggregates of polyQ can siphon essential proteins into amyloid inclusions and that overexpression of these essential proteins can help alleviate toxicity and reduce aggregate size, presumably by rendering fibrils inert by coating their surface^28^. In this way, the different ordered surfaces of E46K fibrils may interact more strongly than wild-type fibrils with certain essential proteins in the cell, for instance those involved in mitochondrial homeostasis, and this may help to explain the greater reduction in mitochondrial activity we observe (Figure 5 b).

### Amyloid polymorphism

The differences in biological activity associated with structural differences between mutant and wild-type fibrils highlight the relevance of amyloid polymorphism to disease. Hereditary mutations may represent one important influence over the formation of different amyloid polymorphs. Here, we learn that the E46K hereditary mutation leads to a new α-syn quasi-polymorph by facilitating a large rearrangement in fibril structure including a re-packed protofilament fold and a new protofilament interface. The polymorphism displayed by E46K is reminiscent of modal polymorphism, first introduced by Caspar and Cohen^29^, whereby identical units have different dispositions in different assemblies (in other words, two fibril structures of the same protein adopt different structures). Although wild-type and E46K fibrils are not strictly modal polymorphs because of the difference in sequence at one amino acid position and hence we refer to them as quasi-polymorphs, it has recently been shown that, under different buffer conditions from our own, the wild-type sequence can indeed form a structure – termed polymorph 2a – similar to the E46K structure determined here^30^. This is consistent with the solvent-facing orientation of sidechain 46 in the E46K structure, thereby compatible with either the negatively charged wild-type glutamate or the positively charged mutant lysine.

The vast difference in structure between the wild-type and E46K fibrils due to a change at a single amino acid position, or a change in fibril growth buffer conditions^30^, highlights the large degree of sensitivity of α-syn fibril assembly to certain interactions. This vulnerability to large-scale rearrangements due to small changes in protein sequence or fibril growth environment reveals that the fibril misfolding landscape for α-syn is flat with many local minima^31^. Here, a single amino acid change, the E46K mutation, shifts the fibril misfolding pathway of α-syn to a different minimum in the folding landscape – the first atomic evidence of a hereditary mutation in an amyloid protein to do so. Energetic analysis reveals that the E46K quasi-polymorph is significantly more stable than the wild-type folds (−0.59 kcal/mol/residue for E46K vs. −0.49 kcal/mol/residue for wild-type rod and −0.30 kcal/mol/residue for wild-type twister, see Table 1). This calculation, along with the fact that the wild-type sequence has been shown to be able to adopt the E46K structure^30^, prompts the question why the wild-type sequence does not always form the more stable E46K structure?

### Kinetic factors that influence amyloid structure

The predominance of one amyloid polymorph over another is depends on stochastic nucleation events and kinetically driven growth processes, permitting less stable polymorphs in a sample to dominate if they form and replicate quicker than more stable polymorphs^32^. We hypothesize that the formation of the E46-K80 salt-bridge observed in all structures of the wild-type rod polymorph determined thus far is an early event in the fibril formation pathway, occurring in the transition state between pre-fibrillar and fibrillary structures, that lowers the energy barrier to forming the rod structure^7,18, 19^. In other words, the early formation of the E46-K80 salt bridge may divert α-syn into a kinetic trap. By constraining the residues connecting Glu46 and Lys80 to adopt a specific conformation that allows Glu46 and Lys80 to maintain their proximity, the E46-K80 salt bridge shifts the misfolding pathway away from the E46K structure and toward the rod structure, thus enforcing the formation of the less stable structure. This salt bridge could be one contributing factor to the observation of the similar kernels formed by residues 50-77 observed in wild-type rod structures^7,18, 19^. The mutation of residue 46 to lysine via the E46K hereditary mutation eliminates the potential to form the E46-K80 salt bridge. This raises the energy barrier to forming the rod and diverts α-syn to a different misfolding pathway, resulting in the more energetically stable E46K structure. These ideas are summarized in Figure 6, which imagines the misfolding landscape of the α-syn rod and E46K quasi-polymorphs (we label the structure on the right the “compact” polymorph in order to avoid confusion as both E46K and other sequences could adopt this fold). This folding landscape is also consistent with our results that E46K α-syn aggregates more slowly than wild-type (Supplementary Figure 5 a) given that the transition state for the wild-type sequence to form the rod structure is predicted to be lower than the transition state for the E46K sequence to form the compact polymorph (Figure 6).

**Figure 6.**
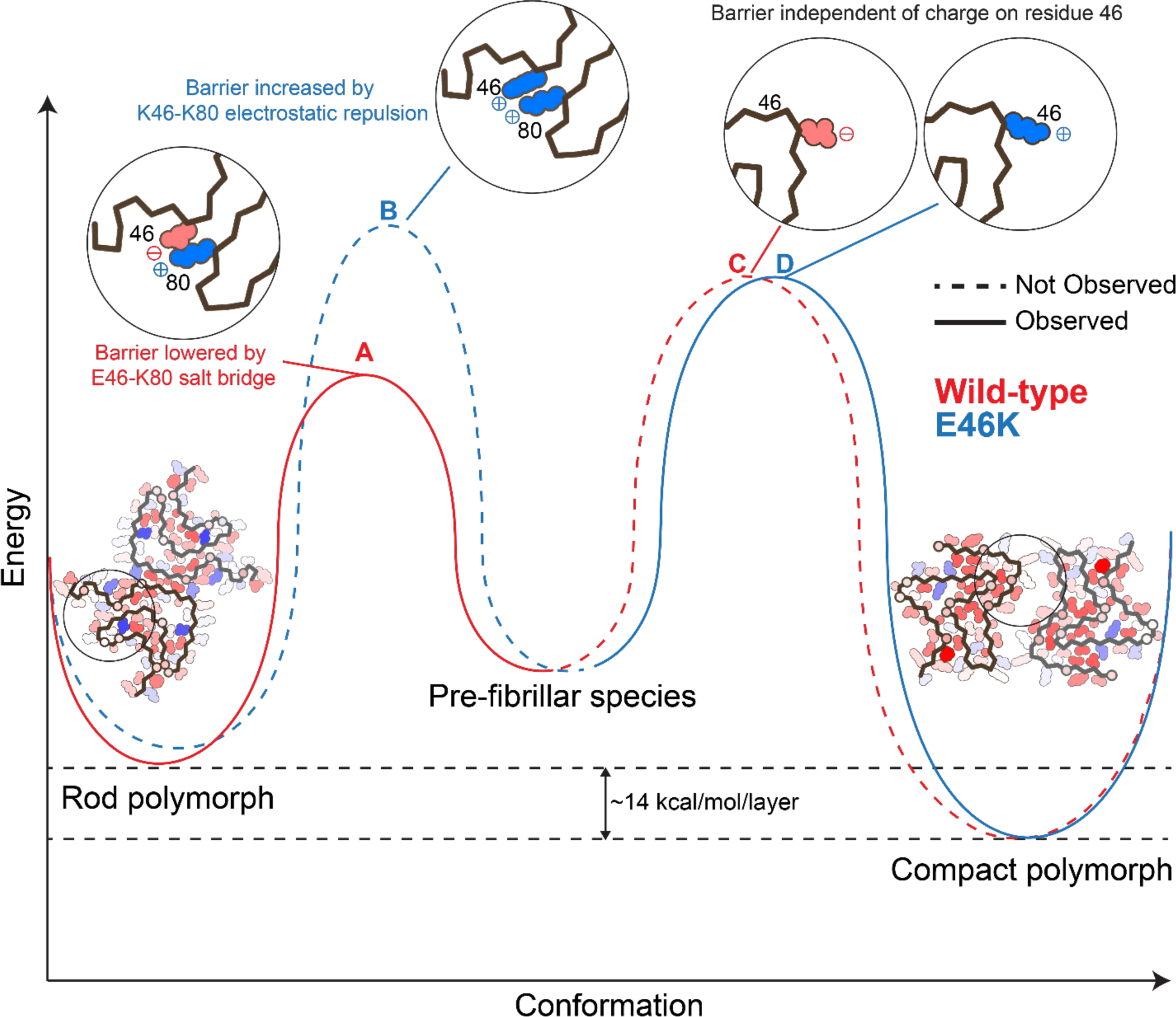
Proposed misfolding landscape of α-syn rod and compact polymorphs The proposed misfolding landscape of α-syn rod and compact (including the E46K quasi-polymorph) polymorphs demonstrates that for the wild-type sequence, it is kinetically more feasible to form the lower energy rod structure due to a transition state whose energy is lowered by the E46-K80 salt-bridge (A). The compact polymorph determined as part of this work is also accessible to the wild-type sequence (C), although it is not observed due to the E46-K80 salt bridge acting as a kinetic trap, diverting the sequence to fold into the rod structure. The E46K mutation thereby serves to raise the transition state energy of forming the rod polymorph via electrostatic repulsion between K46-K80 (B), thus facilitating the formation of the compact polymorph (D).

To date, no other studies of wild-type or mutant α-syn have revealed the twister polymorph we determined in our initial cryo-EM study^7^. It is especially surprising that E46K α-syn does not form the twister structure given that it was predicted that E46K would result in a favorable interaction with E83 in the twister conformation^7^. We speculate that – given the role of kinetics in selecting amyloid polymorphs – the twister polymorph may be the result of a stochastic nucleation event leading to a rare fibril polymorph. Therefore, the twister structure may not be easily reproducible when mutations are added or buffer conditions (influence of fibril growth conditions discussed below) are changed thereby providing an explanation for why we and others have not recapitulated the twister structure in more recent studies.

### Growth conditions that influence amyloid structure

That the wild-type sequence can also adopt the E46K fold adds complexity to the argument that the formation of the E46-K80 salt bridge is an early event in fibril formation that determines the resulting fibril structure^30^. Clearly, there are other factors at play. One such consideration is the role of the fibril growth environment. Guerrero-Ferreira *et al.* note that the juxtaposition of multiple positively charged lysines at positions 43, 45, and 58 seen in the wild-type rod structures may be allowed only in the presence of poly-anionic counter charges such as phosphate^30^. They observe that the removal of phosphate from the buffer conditions disallows this juxtaposition of lysine residues, facilitating the formation of the E46K-like polymorph they term 2a. This finding emphasizes again the sensitivity of α-syn fibril structure to perturbations in sequence or growth environment and that both of these factors play a role in producing new folds. Further, our cross-seeding experiments demonstrating that E46K fibrils can seed wild-type monomer and the fact that the resulting fibrils have different ThT binding ability compared to unseeded wild-type – indicative of a different underlying structure – suggests that the E46-K80 salt-bridge can also be disrupted by templated aggregation with seeds of a different structure (Supplementary Figure 5 a). Together, these findings indicate that the force exerted by the E46-K80 salt bridge is relatively small and can be overcome by different buffer conditions or templated aggregation. However, these small forces have a large influence in selecting the polymorph because they are exerted early in the aggregation pathway.

Interestingly, Guerrero-Ferreira *et al.* also reconstruct a low-resolution density map of E46K α-syn grown in phosphate buffer that resembles the 2a polymorph and our E46K structure determined here^30^. This is identical to the buffer they used previously to grow fibrils of wild-type, C-terminally truncated α-syn that produced a rod polymorph similar to the one we determined^7, 18^. This is similar to the case we describe here in which we grow E46K fibrils using identical buffer conditions as those used to grow fibrils of the wild-type rod polymorph, and the E46K mutation results in a new fold. Taking these results together, this implies that, under identical buffer conditions, E46K acts as a switch to shift the fibril folding pathway, likely through the disruption of the E46-K80 kinetic trap discussed above, thereby unlocking a new, more stable polymorph.

Initial studies examining the effect of the E46K mutation on recombinantly assembled α-syn fibrils further emphasize the importance of fibril growth conditions on selecting polymorphs. Whereas we find that E46K α-syn forms fibrils more slowly than wild-type α-syn (Supplementary Figure 5 a), Choi, *et al.* found that the E46K mutation accelerates α-syn fibril formation^15^. However, the fibrils formed in their study appear to have a different morphology than our E46K fibrils^15^. Specifically, the E46K fibrils prepared under their conditions have a faster twist and greater variation in width along the fibril, indicative of a different underlying molecular structure. This distinction is most likely due to the differences in buffer conditions between our two preparations: Choi *et al.* grew their fibrils in MOPS buffer, whereas we grow our fibrils in water and tetrabutylphosphonium bromide, a fibrillation agent we identified in our initial screening to identify wild-type fibrils suitable for cryo-EM structure determination^7^. The observed differences in fibril morphology and aggregation kinetics again highlight the sensitivity of α-syn fibrillation to growth conditions and hints at the ability of E46K α-syn to form still further polymorphs not determined in this work.

### The folding landscape of amyloid proteins

As mentioned above, amyloid polymorphism is likely due to a flat protein folding landscape in which many local minima exist. This landscape contrasts with that of most proteins whose sequences have evolved to encode one structure with the lowest free energy in a funnel-shaped protein folding landscape^31, 33^. The prodigious polymorphism observed not only in α-syn structures determined in our work but in other amyloid structures hints that evolution has not played a role in specifying pathogenic amyloid structures. Therefore, many factors, such as concentration and intra-chain interactions, may guide the sequence to forming a non-evolved structure. The E46-K80 salt bridge may represent one of these intra-chain interactions that diverts wild-type α-syn into a local energy minimum (i.e., a kinetic trap), and the E46K hereditary mutation unlocks this constraint, allowing a more stable structure to form.

The release of local constraints leading to lower free energy structures could be a mechanism by which other hereditary mutations operate as well. This hypothesis is in line with our previous studies showing that pathogenic mutations and post-translational modifications can lead to more stable amyloid assemblies^34–36^. That work was done on shorter peptide segments however and may not have captured all the interactions present in a full-length amyloid fibril. Therefore, future work is needed to compare structures of full-length wild-type and hereditary mutant or post-translationally modified amyloid fibrils to identify the possible mechanisms by which hereditary mutations and PTM’s can alter amyloid protein pathogenicity. We have previously shown that the H50Q hereditary mutation alters α-syn’s protofilament assembly, resulting in more pathogenic fibrils – a first step in understanding the role of hereditary mutations in fibril structure and activity^37^. Recently, a pair of studies demonstrated that N-terminally acetylated α-syn has different ThT binding ability and different seeding properties compared to non-acetylated α-syn although the structures of non-acetylated and acetylated α-syn fibril cores remain largely similar^20, 25^. Further, ssNMR studies of α-syn hereditary mutants A30P, which lies outside the fibril core of α-syn structures determined to date, and A53T, which lies at the protofilament interface of wild-type rod polymorphs, suggest only local perturbations compared to the wild-type fibril structure^14, 38^. Also, hereditary mutations in transthyretin have been discovered that serve to de-stabilize transthyretin’s native fold thereby promoting fibril formation^39^. Together, these data indicate that other modes of mutationally- or PTM-encoded pathogenicity may exist including effects localized to regions outside the fibril core or effects on the monomeric protein structure. Indeed, PTM’s such as phosphorylation, oxidative stress and truncation have varying effects on α-syn aggregation and toxicity^40^.

## Conclusion

In summary, we have determined a 2.5 Å resolution reconstruction of recombinantly assembled E46K α-syn fibrils that provides the first atomic structure of this hereditary mutation initially discovered in a family with a clinical diagnosis of parkinsonism and Lewy Body Dementia^13^. The fibril structure of E46K α-syn greatly differs from and has a lower free energy than wild-type structures, and we attempt to use the structure to rationalize its higher seeding capacity and mitochondrial impairment compared to wild-type. We posit that due to fibril α-syn’s unevolved nature, the E46-K80 salt bridge in wild-type fibrils represents a local constraint that prevents the formation of the lower free energy fibril fold; E46K alleviates this constraint allowing re-folding to a more stable structure. The release of local constraints to allow re-packing into more stable, pathogenic fibrils may be a mechanism by which other hereditary mutations operate in α-syn and other amyloid proteins.

## Acknowledgments

We thank H. Zhou for use of Electron Imaging Center for Nanomachines (EICN) resources and P. Ge for assistance in cryo-EM data collection. We acknowledge the use of instruments at the EICN supported by NIH (1S10RR23057 and 1S10OD018111), NSF (DBI-1338135), and CNSI at UCLA. The authors acknowledge NIH AG 060149, NIH AG 054022, NIH AG061847, and DOE DE-FC02-02ER63421 for support. D.R.B. was supported by the National Science Foundation Graduate Research Fellowship Program.

## Author Contributions

D.R.B. and B.L. designed experiments and performed data analysis. B.L. and C.S. expressed and purified the α-syn protein. B.L. grew fibrils of α-syn and performed biochemical experiments. D.R.B. and B.L prepared cryo-EM samples and performed cryo-EM data collection. B.L. and W.F. selected filaments from cryo-EM images. D.R.B. performed cryo-EM data processing and built the atomic models. M.R.S. wrote the software for and D.R.B. carried out solvation energy calculations. M.P.H wrote the software for and D.R.B. carried out the pairwise interaction analysis. All authors analyzed the results and D.R.B wrote the manuscript with input from all authors. L.J. and D.S.E. supervised and guided the project.

## Data Availability Statement

All structural data have been deposited into the wwPDB and EMDB with the following accession codes: PDB 6UFR and EMDB EMD-20759. All other data are available from the authors upon reasonable request.

## Supporting Information

### Methods

#### Protein purification

Full-length α-syn wild-type and E46K mutant proteins were expressed and purified according to a published protocol^1^. Transformed bacteria were induced at an OD600 of ∼0.6 with 1 mM IPTG for 6 h at 30°C. The bacteria were then lysed with a probe sonicator for 10 minutes in an iced water bath. After centrifugation, the soluble fraction was heated in boiling water for 10 minutes and then titrated with HCl to pH 4.5 to remove the pellet. After adjusting to neutral pH, the protein was dialyzed overnight against Q Column loading buffer (20 mM Tris-HCl, pH 8.0). On the next day, the protein was loaded onto a HiPrep Q 16/10 column and eluted using elution buffer (20 mM Tris-HCl, 1M NaCl, pH 8.0). The eluent was concentrated using Amicon Ultra-15 centrifugal filters (Millipore Sigma, 10 NMWL) to ∼5 mL. The concentrated sample was further purified with size-exclusion chromatography through a HiPrep Sephacryl S-75 HR column in 20 mM Tris, pH 8.0. The purified protein was dialyzed against water, concentrated to 3 mg/ml, and stored at 4 °C. The concentration of the protein was determined using the Pierce™ BCA Protein Assay Kit (cat. No. 23225, Thermo Fisher Scientific).

#### Fibril preparation and optimization

Both wild-type and E46K fibrils were grown under the same condition: 300 µM purified monomers, 15mM tetrabutylphosphonium bromide, shaking at 37°C for 2 weeks.

#### Fibril seeding aggregation in cells

We performed the biosensor cell seeding assay based on a previously published protocol^2^. Briefly, the assay works as follows: exogenous, un-labeled fibrils are transfected into HEK293T cells expressing α-syn-A53T-YFP. Seeded aggregation of endogenously expressed α-syn-A53T-YFP is monitored by formation of fluorescent puncta. The puncta represent intracellular aggregation of α-syn-A53T-YFP as a result of seeding by exogenous E46K or WILD-TYPE fibrils.

Human embryonic kidney FRET Biosensor HEK293T cells expressing full-length α-syn containing the hereditary A53T mutation were grown in DMEM (4mM L-glutamine and 25mM D-glucose) supplemented with 10% FBS, 1% penicillin/streptomycin. Trypsin-treated HEK293T cells were harvested, seeded on flat 96-well plates at a concentration of 4×104 cells/well in 200 µL culture medium per well and incubated in 5% CO2 at 37°C for 18 hours.

α-syn fibrils were prepared by diluting with Opti-MEM™ (Life Technologies; Carlsbad CA) and sonicating in a water bath sonicator for 10 minutes. Fibril concentration was determined as monomer-equivalent concentration. The fibril samples were then mixed with Lipofectamine™ 2000 (Thermo Fisher Scientific) and incubated for 15 minutes and then added to the cells. The actual volume of Lipofectamine™ 2000 was calculated based on the dose of 1 µL per well. After 48 hours of transfection, the cells were trypsinized, transferred to a 96-well round-bottom plate and resuspended in 200 µL chilled flow cytometry buffer (HBSS, 1% FBS, and 1 mM EDTA) containing 2% paraformaldehyde. The plate was sealed with Parafilm and stored at 4 °C for imaging. Fluorescent images were processed in ImageJ to count number of seeded cells.

#### MTT mitochondrial activity assay

The addition of sonicated fibrils to Nerve Growild-typeh Factor-differentiated PC12 cells is a well-established assay to measure cytotoxicity of amyloid fibrils^1, 3–5^. Use of this neuron-like cell line allows us to obtain a biologically relevant assay for cytotoxicity. For our MTT mitochondrial activity assay, the protocol was adapted from the Provost and Wallert laboratories and was performed in an identical manner to our previous work^5, 6^. Thiazolyl blue tetrazolium bromide for the MTT cell toxicity assay was purchased from Millipore Sigma (M2128-1G; Burlington, MA). PC12 cells were plated in 96-well plates at 10,000 cells/well in DMEM (Dulbecco’s modification of Eagle’s medium; 5% fetal bovine serum [FBS], 5% heat-inactivated horse serum, 1% penicillin/streptomycin and 150 ng/mL nerve growild-typeh factor 2.5S (Thermo Fisher Scientific). The cells were incubated for 2 days in an incubator with 5% CO2 at 37°C. The cells were treated with different concentrations of monomer-equivalent α-syn fibrils (200 nM, 500 nM, 1000 nM, and 2000 nM). After 18 hours of incubation, 20 µl of 5 mg/ml MTT was added to every well and the plate was returned to the incubator for 3.5 hours. With the presence of MTT, the experiment was conducted in a laminar flow hood with the lights off and the plate was wrapped in aluminum foil. The media was then removed with an aspirator and the remaining formazan crystals in each well were dissolved with 100 µL of 100% DMSO. Absorbance was measured at 570 nm to determine the MTT signal and at 630 nm to determine background. The data were normalized to those from cells treated with 1% SDS to obtain a value of 0%, and to those from cells treated with PBS to obtain a value of 100%.

#### In Vitro Aggregation and Seeding Assay

α-syn wild-type or E46K monomers (100 µM) were mixed with 60 µM thioflavin-T (ThT) and transferred into a 96-well plate. The signal was monitored using the FLUOstar Omega Microplate Reader (BMG Labtech, 37 °C with 600 r.p.m. double orbital shaking, ex. 440 nm, em. 490 nm). For the seeding groups, preformed wild-type or E46K fibrils (10 µM) after 10 minutes of water-bath sonication were added to the wild-type α-syn monomers immediately before beginning the aggregation assay.

#### SDS Stability Assay

The wild-type and E46K aggregated α-syn samples at the end of the ThT assay were treated with addition of 10% SDS to reach SDS final concentration of 0.5%. The ThT signal is measured after 5 minutes of incubation at 37 °C with 600 r.p.m. double orbital shaking. The addition of SDS and ThT measurement is repeated with increments of 0.5% to a final SDS concentration of 3.5%. The initial ThT signals at 0% SDS were used for normalization.

#### Cryo-EM data collection and processing

2 µl of fibril solution was applied to a baked and glow-discharged Quantifoil 1.2/1.3 electron microscope grid and plunge-frozen into liquid ethane using a Vitrobot Mark IV (FEI). Data were collected on a Titan Krios (FEI) microscope equipped with a Gatan Quantum LS/K2 Summit direct electron detection camera (operated with 300 kV acceleration voltage and slit width of 20 eV). Counting mode movies were collected on a Gatan K2 Summit direct electron detector with a nominal physical pixel size of 0.843 Å/pixel with a dose per frame 1.2 e-/Å^2^. A total of 30 frames with a frame rate of 5 Hz were taken for each movie resulting in a final dose of 36 e-/Å^2^ per image. Automated data collection was driven by the SerialEM automation software package with image shift induced-beam tilt correction^7^.

Micrographs containing crystalline ice were used to estimate the anisotropic magnification distortion using mag_distortion_estimate^8^. CTF estimation was performed using CTFFIND 4.1.8 on movie stacks with a grouping of 3 frames and correction for anisotropic magnification distortion^9^. Unblur^10^ was used to correct beam-induced motion with dose weighting and anisotropic magnification correction, resulting in a physical pixel size of 0.838 Å/pixel.

All particle picking was performed manually using EMAN2 e2helixboxer.py^11^. All fibril particles were first extracted using 1024 pixel box sizes and a 10% inter-box distance then subjected to 2D class averaging. 2D class averages reveal that E46K α-syn forms fibrils of a single morphology with a pitch of ∼800 Å (Supplementary Figure 2b). We next extracted all fibrils with a 686 pixel box size and 10% inter-box distance, and again performed 2D class averaging. 2D class averaging of 686 pixel boxes resulted in clear separation of beta-strands along the length of the fibril. 2D class averages and their corresponding simulated diffraction patterns together indicate that the helical rise is ∼4.8 Å with a C_n_ rotational symmetry due to the presence of a meridional reflection (Supplementary Figure 2b). Due to the two-fold mirror symmetry present in the 2D class averages, we reasoned that the fibril had a C_2_ rotational symmetry. Using a calculated helical twist of 178.92° (given pitch of 800 Å and rise of 4.8 Å) and C_2_ rotational symmetry, we carried out 3D class averaging with a single class and a featureless cylinder created by relion_helical_toolbox as an initial model^12^. The featureless cylinder was refined to a reasonable model where separation of beta-sheets in the x-y plane of the fibril could be visualized. This model was then used to separate good and bad particles with a 3D class averaging job with three classes. Fibrils contributing to the best class in the previous 3D classification were re-extracted with a 400 pixel box size, 10% inter-box distance, and phase flipped for subsequent classification and high resolution refinement. An additional 3D class averaging job was performed and a final subset of 114,260 helical segments were selected for gold-standard auto-refinement in RELION. Refinement yielded a final 2.5 Å reconstruction (Supplementary Figure 2c). We sharpened the map using phenix.auto_sharpen with a sharpening factor of −140 Å^2^ and a resolution cutoff of 2.5 Å^13^.

#### Atomic model building

We used phenix.map_to_model with an input sequence corresponding to E46K α-syn to build an initial model^13^. Phenix.map_to_model correctly built a segment of the N-terminus of the fibril core and we manually built the rest of the structure into the density map in COOT^14^. We generated a 5-layer model to maintain local contacts between chains in the fibril during structure refinement. We performed automated structure refinement using phenix.real_space_refine^13^. We employed hydrogen bond distance and angle restraints for backbone atoms participating in β-sheets and side chain hydrogen bonds during automated refinements. We performed comprehensive structure validation of all our final models in Phenix.

#### Energetic calculation

The standard free energy of stabilization of a given amyloid chain is computed as the difference in atomic solvation energy of a pseudo-extended, solvated chain and the folded chain in the center of five layers of the known structure of a protofilament^15^. The atomic solvation parameters are those of Eisenberg *et al*. (1989), with additional terms to describe the entropy change of sidechains on folding, as calculated by Koehl and Delarue (1994), scaled by the percentage of side chain surface area buried^16, 17^.

---

**Supplementary Figure 1.**
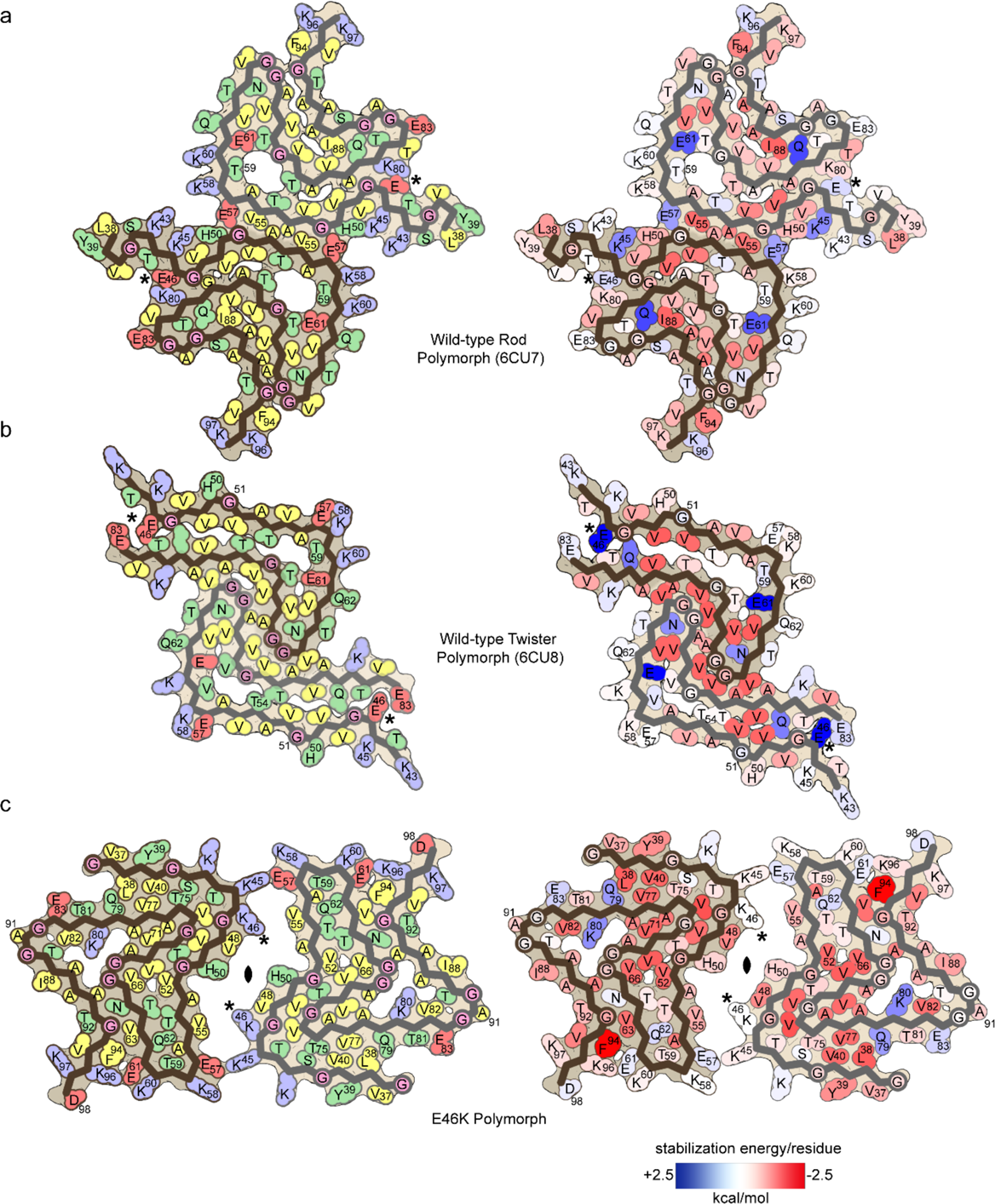
Schematic representation and free energy of stabilization maps for the wild-type and E46K polymorphs a-c) (left) Schematic representation of fibril structures with amino acid side chains colored as follows: hydrophobic (yellow), negatively charged (red), positively charged (blue), polar, uncharged (green), and glycine (pink). (right) Solvation energy maps of fibril structures. The stabilizing residues are red; the de-stabilizing residues are blue.

**Supplementary Figure 2.**
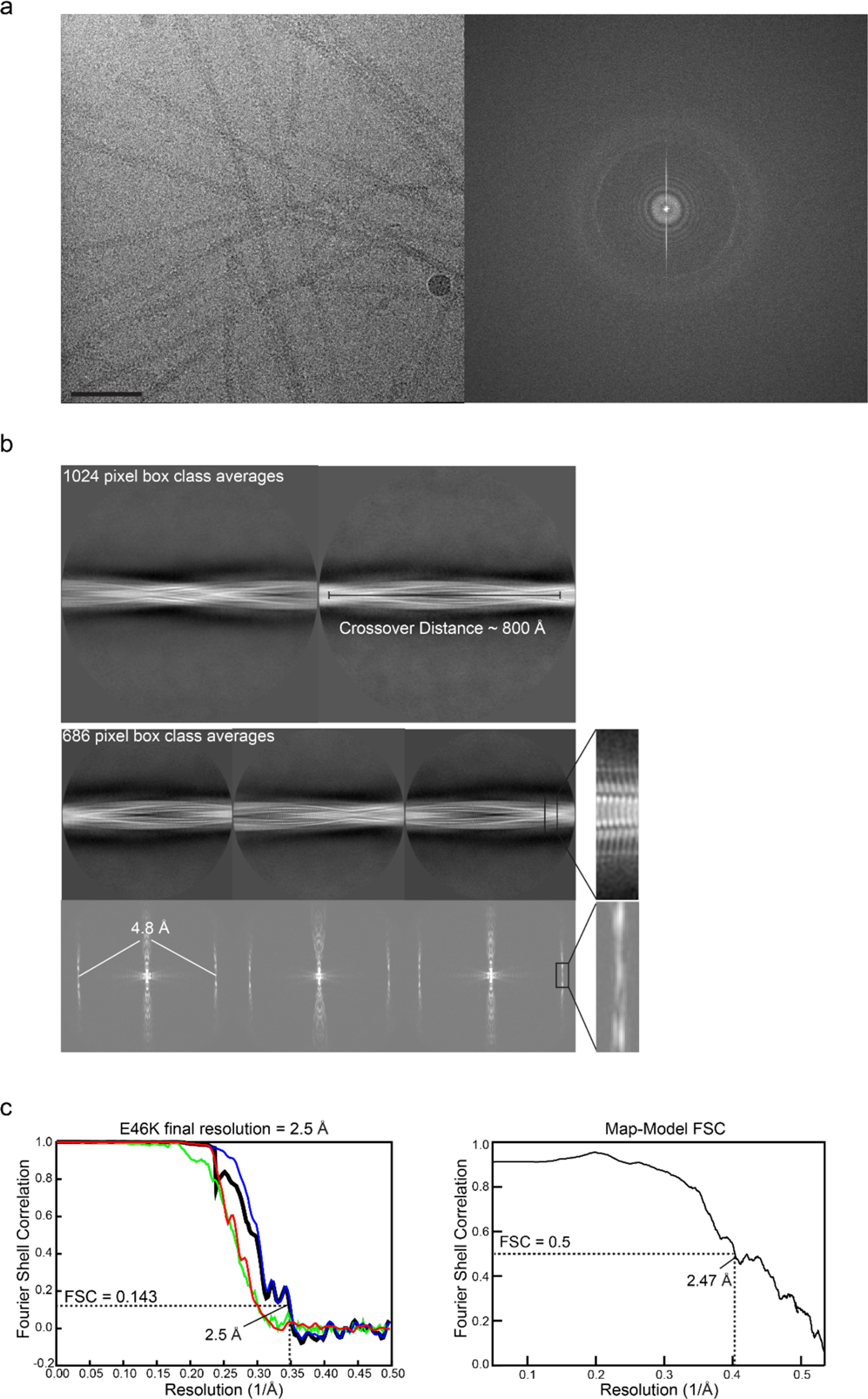
Cryo-EM Data Collection and Processing a) Representative cryo-EM image and power spectrum. Scale bar 50 nm. b) 1024 pixel box class averages reveal a helical pitch of ∼800 Å. 686 pixel box class averages and corresponding simulated diffraction patterns reveal a rise of ∼4.8 Å and a C_2_ rotational symmetry. c) Gold-standard half-map-half-map FSC and map-to-model FSC.

**Supplementary Figure 3.**
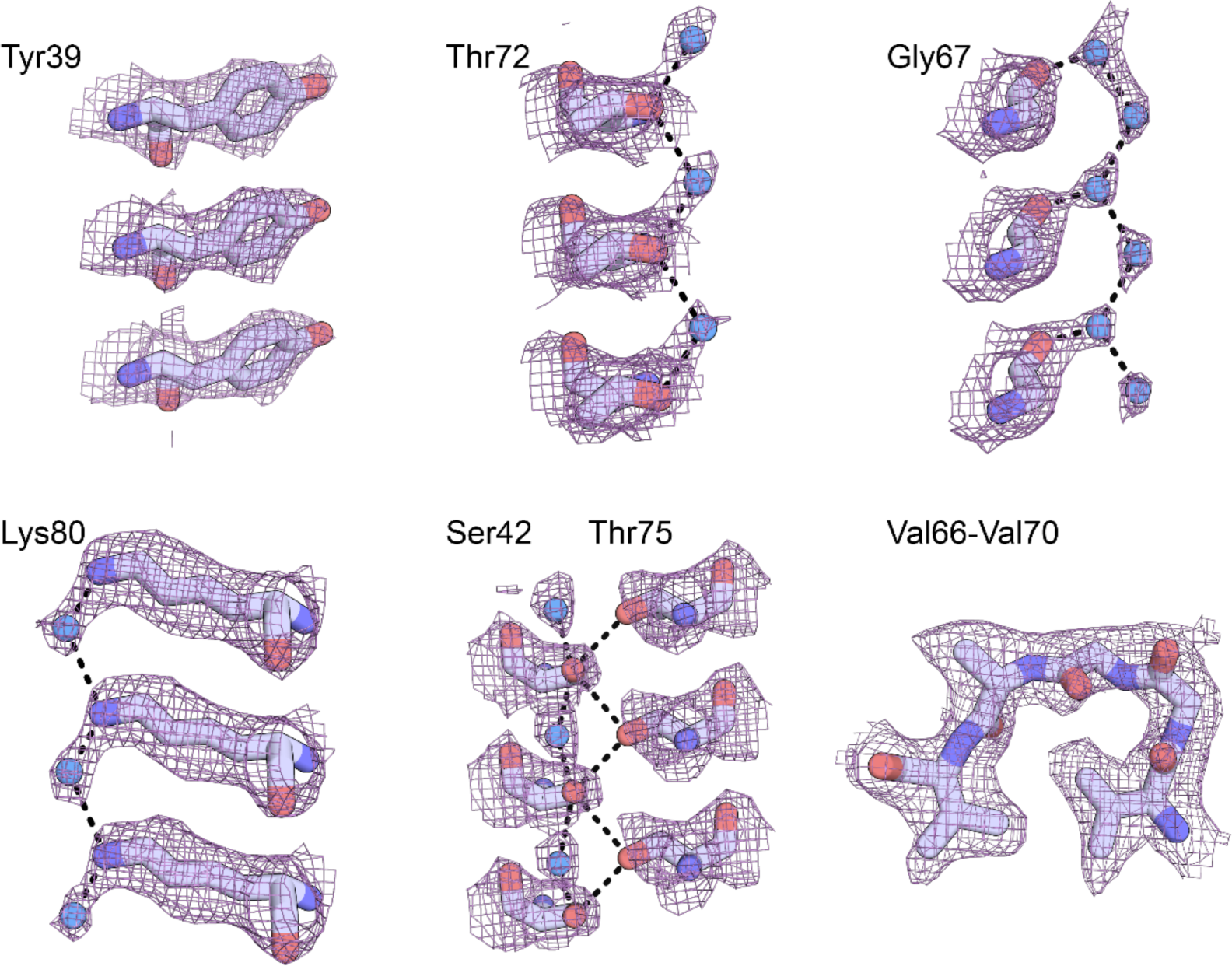
Atomic model and density map highlight resolution of reconstruction and hydrogen bonding networks.

**Supplementary Figure 4.**
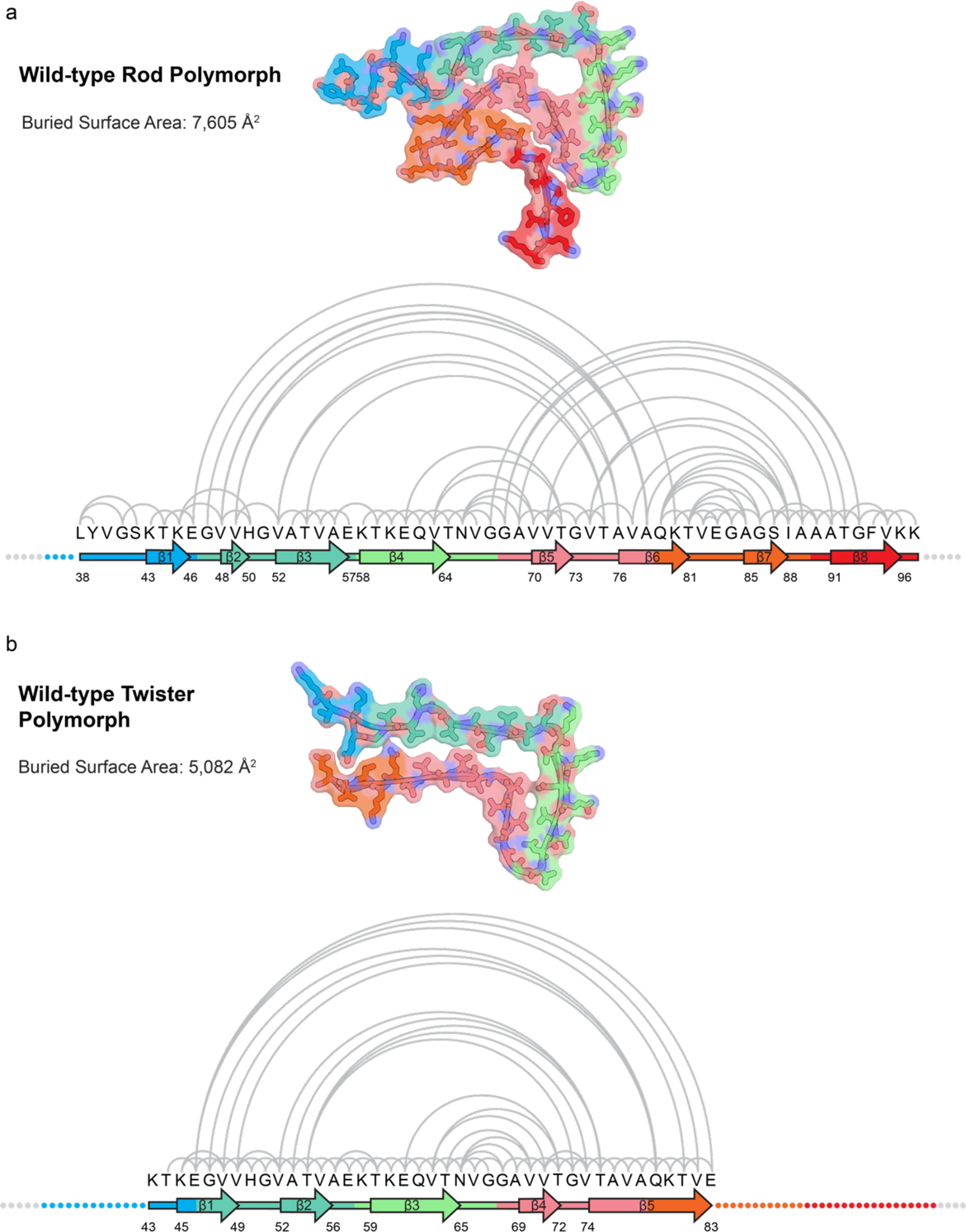
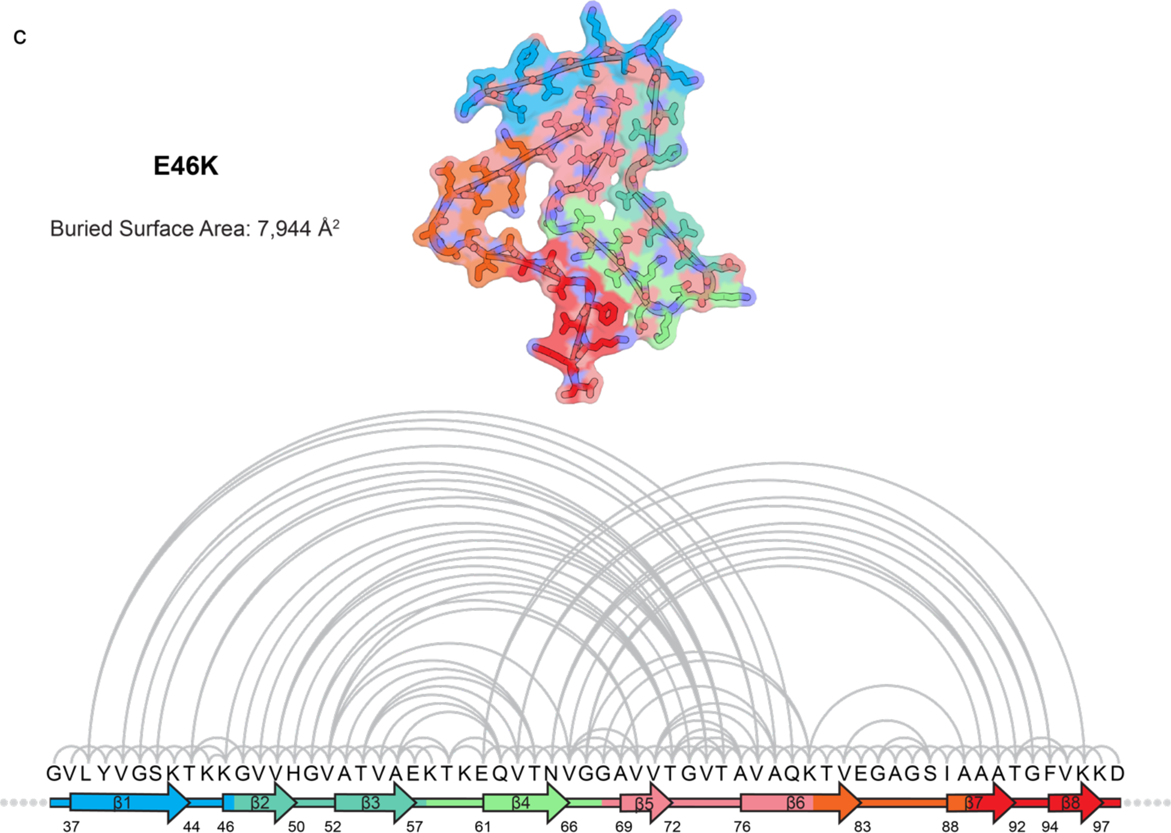
Pairwise interaction analysis of wild-type and E46K protofilament folds a-c) Protofilament fold and calculated buried surface area (top) and fibril core primary and secondary structure with pairs of interacting residues connected by half-circles (bottom) for a) wild-type rod, b) wild-type twister, and c) E46K polymorphs. E46K polymorph has a different set of interacting residues highlighting the difference in protofilament fold compared to wild-type. The E46K polymorph also has a larger number of interactions, which is reflected in its higher buried surface area.

**Supplementary Figure 5.**
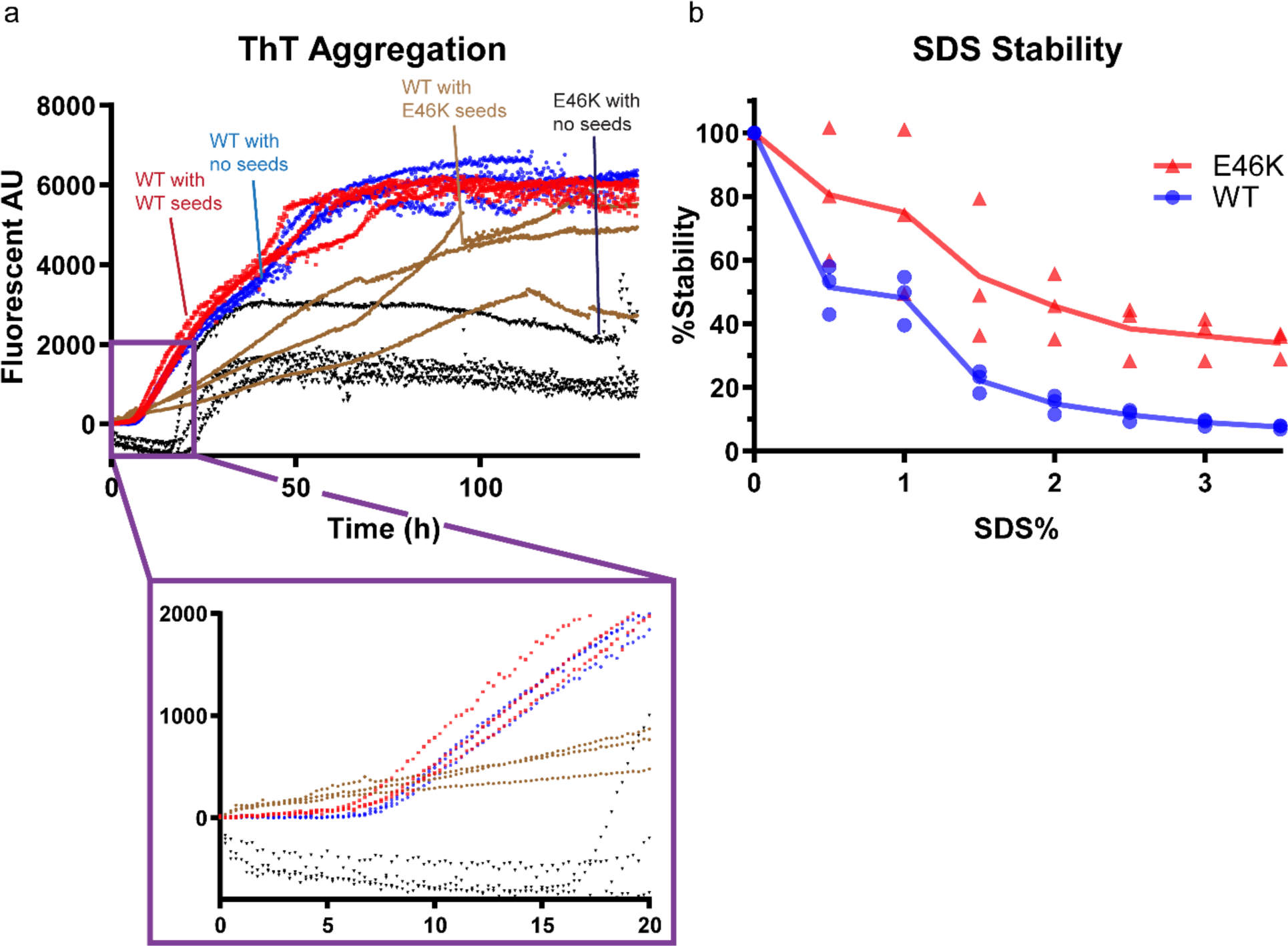
Cross-seeding of wild-type α-syn by E46K fibrils and SDS fibril stability assay a) Seeding of wild-type α-syn by wild-type fibrils results in a modestly reduced lag-time while seeding by E46K fibrils eliminates the lag phase. In addition, both unseeded and self-seeded wild-type α-syn have similar ThT binding ability shown by their similar ThT aggregation curves; on the other hand, wild-type α-syn seeded by E46K fibrils has a different ThT fluorescence intensity, indicating a different underlying structure. Unseeded wild-type α-syn has a shorter lag phase and higher max ThT signal than E46K α-syn. Breaks in the ThT curves originate from the microplate-reader being interrupted and re-started to allow other experiments to be performed in separate wells in the same microplate. Due to normalizing the initial reading to zero, the E46K lag phase dips below zero fluorescence AU. The reason for this slight dip after the initial ThT reading is unknown; however, the classical nucleation-elongation sigmoidal growth curve is still demonstrated for E46K α-syn. b) E46K and wild-type fibrils were heated to 37 °C and incubated with varying concentrations of SDS. E46K fibrils are more resistant to SDS than wild-type fibrils. Individual triplicate measurements are shown and the plotted line represents the average of the triplicates.

